# Female-predominant anti-CD4 IgG autoantibody production and its correlations with plasma levels of progesterone, microbial translocation, and blunted immune reconstitution in HIV-infected patients on suppressive ART

**DOI:** 10.64898/2025.12.03.689730

**Authors:** Zhuang Wan, Zhenwu Luo, John E. McKinnon, Alicia Hartley, Richard W Price, Magnus Gisslen, Wei Jiang

## Abstract

Autoimmunity contributes to HIV immunopathogenesis even in the absence of overt autoimmune disease. We previously showed that anti-CD4 autoantibodies from people with HIV (PWH) on suppressive antiretroviral therapy (ART) can mediate cytotoxicity against CD4+ T cells, implicating a role in impaired immune reconstitution. Despite viral suppression, many PWH with poor CD4 recovery exhibit chronic immune activation, microbial translocation, and dysregulated humoral immunity. Here, we identify a female-predominant elevation of plasma anti-CD4 IgG autoantibodies in aviremic PWH receiving ART. Across two independent cohorts, HIV-positive females, but not males, displayed significantly higher anti-CD4 IgG, predominantly IgG1, compared with HIV-negative controls, without parallel increases in anti-CD4 IgA or IgM. This sex-specific pattern was unique to anti-CD4 IgG and was not observed for anti-CD8 IgG, anti-double-stranded DNA IgG, or anti-nuclear antigen IgG; these control autoantibodies correlated with one another but not with anti-CD4 IgG. Elevated anti-CD4 IgG levels were associated with lower plasma progesterone levels and reduced absolute CD4+ T-cell counts. Markers of microbial translocation, soluble CD14 (sCD14), lipopolysaccharide-binding protein (LBP), and lipopolysaccharide (LPS), were also selectively increased in HIV-positive females, with sCD14 and LBP showing significant or borderline associations with anti-CD4 IgG. Together, these findings identify anti-CD4 IgG as a sex-dimorphic autoimmune signature in treated HIV infection, linked to progesterone levels, persistent microbial translocation, and incomplete immune recovery. This work highlights an under-recognized intersection of sex, mucosal barrier dysfunction, and autoimmunity in HIV pathogenesis and suggests potential therapeutic targets to improve immune reconstitution in women.

## INTRODUCTION

In 2017, we reported the first evidence that autoimmunity can drive HIV immunopathogenesis without overt autoimmune disease (1). This paradigm has since been widely embraced and expanded, shaping current understanding of autoimmunity-mediated immune dysregulation in other infectious disorders such as COVID-19 (2). Notably, about 20-25% of people with HIV (PWH) receiving antiretroviral therapy (ART) fail to fully reconstitute CD4+ T-cell counts despite achieving plasma viral suppression, resulting in elevated risks of comorbidities and mortality (3). We previously demonstrated that anti-CD4 autoantibodies drive CD4+ T-cell death through antibody-dependent cellular cytotoxicity (ADCC), thereby contributing to incomplete immune recovery on ART (1). The role of anti-CD4 IgG in CD4+ T-cell depletion has also been described by Lisco et al. (4) and remains an active area of investigation with heterogeneous findings (5). However, the prevalence, specificity, and clinical correlates of anti-CD4 autoantibodies in PWH on suppressive ART remain poorly defined.

Sex differences profoundly influence immune responses. Females generally mount stronger humoral and cellular immunity but are at higher risk for autoimmune diseases (6). Our previous study on PWH on suppressive ART revealed that females had increased influenza-specific antibody avidity relative to males, but similar plasma levels of influenza-specific binding antibodies and neutralizing antibodies (7). Furthermore, higher anti-CD4 autoantibody levels have been shown in HIV-infected women versus men during pre-ART (8), and women exhibit lower viral loads pre-ART but faster CD4+ T cell decline post-infection and differential responses to therapy (9). Whether these disparities extend to autoantibody profiles in treated HIV remains unknown.

Chronic immune activation and inflammation persist in PWH despite suppressive ART, contributing to non-AIDS comorbidities including cardiovascular disease, neurocognitive decline, and metabolic disorders (10). A hallmark of this dysregulated state is microbial translocation from the gastrointestinal tract, as evidenced by elevated plasma levels of lipopolysaccharide (LPS), LPS-binding protein (LBP), and soluble CD14 (sCD14), a surrogate marker of monocyte activation (11, 12).

In the current study, we comprehensively profiled anti-CD4 autoantibodies in sex differences, immunoglobulin subclasses, and microbial translocation. We found that anti-CD4 IgG uniquely characterizes female-predominant autoantibodies in treated HIV disease and correlates with microbial translocation and poor immune reconstitution with suppressive ART.

## MATERIALS AND METHODS

### Study subjects

The prospective study enrolled 42 study participants at the Medical University of South Carolina (MUSC), Charleston, SC, USA, including 28 aviremic PWH on stable ART (17 HIV+ male, 9 HIV+ female) and 16 HIV-seronegative controls (5 non-HIV males, 11 non-HIV females) in the HIV cohort. The second HIV cohort (13), a retrospective study including 16 non-HIV males, 22 HIV+ males on suppressive ART, and 4 HIV+ females on suppressive ART, was from Sahlgrenska University Hospital, Gothenburg, Sweden, and San Francisco General Hospital, University of California, San Francisco (UCSF), USA. The local Institutional Review Board approved the study. Informed consent was obtained from all participants. The inclusion criteria in the MUSC cohort are: age ≥18 years; for PWH, confirmed HIV-1 infection, stable ART for at least one year, HIV RNA < 40 copies/mL; and willingness to provide blood samples. The exclusion criteria in the MUSC cohort are: pregnancy, breastfeeding, hemoglobin <6.5 g/dL, acute illness, immunomodulatory therapy (including >5 mg/day prednisone) within 120 days, or investigator-determined unsuitability. The second cohort has been described in our previously published study (13). The clinical characteristics of the MUSC cohort are shown in Tables 1 and the second cohort are shown previously (13).

**Table 1.** Clinical characteristics of the MUSC cohort.

| **Characteristics** | **Non-HIV-M** | **Non-HIV-F** | **HIV-M** | **HIV-F** | **P value** |
| --- | --- | --- | --- | --- | --- |
| N (M/F) | 5 | 11 | 17 | 9 |  |
| Age (mean y, SD) | 34.4 (9.3) | 45.5 (10.5) | 39.2 (12.3) | 48.2 (9.0) | NS |
| Race (n, %) |  |  |  |  | NS |
| White | 1 (20) | 5 (45) | 9 (53) | 2 (22) |  |
| Black | 4 (80) | 6 (55) | 8 (47) | 7 (78) |  |
| Others | 0 (0) | 0 (0) | 0 (0) | 0 (0) |  |
| CD4 count, cells/uL | 496 (216) | 813 (370) | 624 (319) | 541 (283) | NS |
| ART duration, year |  |  | 4.3 (1.9) | 5.4 (1.2) | NS |
| Nadir CD4 count |  |  | 339 (177) | 331 (350) | NS |

### Sample Collection

Blood was collected in EDTA and heparin tubes. Plasma was isolated by centrifugation (800 × *g*, 10 min), aliquoted, and stored at −80°C. Peripheral blood mononuclear cells (PBMCs) were isolated by Ficoll-Hypaque density gradient (GE Healthcare, Chicago, Illinois), viably cryopreserved in 10% DMSO and 90% FBS, and stored in liquid nitrogen.

### ELISA for soluble CD4 (sCD4), anti-CD4, anti-CD8, anti-double stranded DNA (dsDNA), and anti-nuclear antigen (ANA) IgGs

96-well plates (high-binding plate, Santa Cruz Technology, Dallas, Texas) were coated overnight at 4°C with recombinant human CD4 (5 μg/mL, Sino Biological), CD8 (5 μg/mL), dsDNA (10 μg/mL), anti-CD4 antibody (5 μg/mL), or nuclear extract (for ANA) in PBS. Plates were blocked with 1% BSA/PBS, incubated with plasma (1:50-1:200), and detected with HRP-conjugated anti-human IgG, IgG1-4, IgA, or IgM (Southern Biotech, Homewood, Alabama). 3,3′, 5,5′ tetramethylbenzidine dihydrochloride (TMB) substrate was used; reactions stopped with 1N H_2_SO_4_. Absorbance was read at 450 nm (OD_450_). Standard curves (purified human IgG) quantified anti-CD4 IgG and anti-dsDNA IgG (ng/mL); others were reported as OD_450_. The assessment of sCD4 levels was described previously (1).

### Autoantibody array for anti-CD4 IgGs

As described previously (13), plasma anti-CD4 IgG levels were evaluated using an autoantibody array (Microarray and Immune Phenotyping Core Facility at the University of Texas Southwestern Medical Center) for the second HIV cohort. The averaged net fluorescent intensity (MFI) of each human CD4 protein autoantigen was normalized to internal controls, and scaling and centering were performed after log10 transformation using the following formula: (x – mean[x])/sd(x).

### Progesterone, 17β-estradiol, and testosterone

The plasma levels of the three sex hormones were measured using ELISA kits (Abcam, Waltham, Massachusetts) according to the manufacturer’s instructions.

### sCD14, LBP, and LPS

As described in our previous study (14), commercial ELISA kits were used per manufacturer: sCD14 (R&D Systems, Minneapolis, MN, USA), LBP (Hycult Biotech, Uden, Netherlands), LPS (ThermoFisher, Florence, SC, USA), sCD4 (BioLegend, San Diego, California, USA).

### Statistical Analysis

Data expressed as median (IQR). Group comparisons were analyzed using Kruskal-Wallis with Dunn’s post-hoc or one-way ANOVA with Tukey’s method for multiple comparisons. Sex-stratified differences were analyzed using Mann-Whitney U. Correlations were analyzed using Spearman’s rank. *p* ≤ 0.05 is considered significant. Analyses performed in GraphPad Prism 9 and SPSS v16.

## RESULTS

### Female-predominant anti-CD4 IgG in PWH on suppressive ART

The medians of plasma anti-CD4 IgG levels in the MUSC cohort were 612 ng/mL (IQR 420–890), 198 ng/mL (IQR 110–320), 82 ng/mL (IQR 45–135), and 68 ng/mL (IQR 40–110) in HIV+ females (HIV-F), HIV+ males (HIV-M), control females (non-HIV-F), and control males (non-HIV-M), respectively (Figure 1A; ANOVA *p* < 0.05). HIV+ females, but not HIV+ males, exhibited increased anti-CD4 IgG compared to controls (*p* < 0.05). In the second retrospective cohort using microarray, a different method, the median MFIs of plasma anti-CD4 IgG were 7917 (IQR 845–14863), 430 (IQR 259-873), and 437 (IQR 312-533) in HIV+ females (HIV-F), HIV+ males (HIV-M), and control males (non-HIV-M), respectively (Figure 1A; ANOVA *p* < 0.05). Anti-CD4 IgG levels were significantly elevated in HIV+ female groups compared to controls. Subclass analysis revealed IgG1 as the dominant isotype in HIV-F compared to non-HIV-F (OD_450_ 0.48 vs. 0.22, *p* = 0.016, Mann-Whitney; Figure 1B). IgG2–4 showed no differences. Anti-CD4 IgA and IgM were similar across all groups (Supplemental Figure 1).

**Figure 1.**
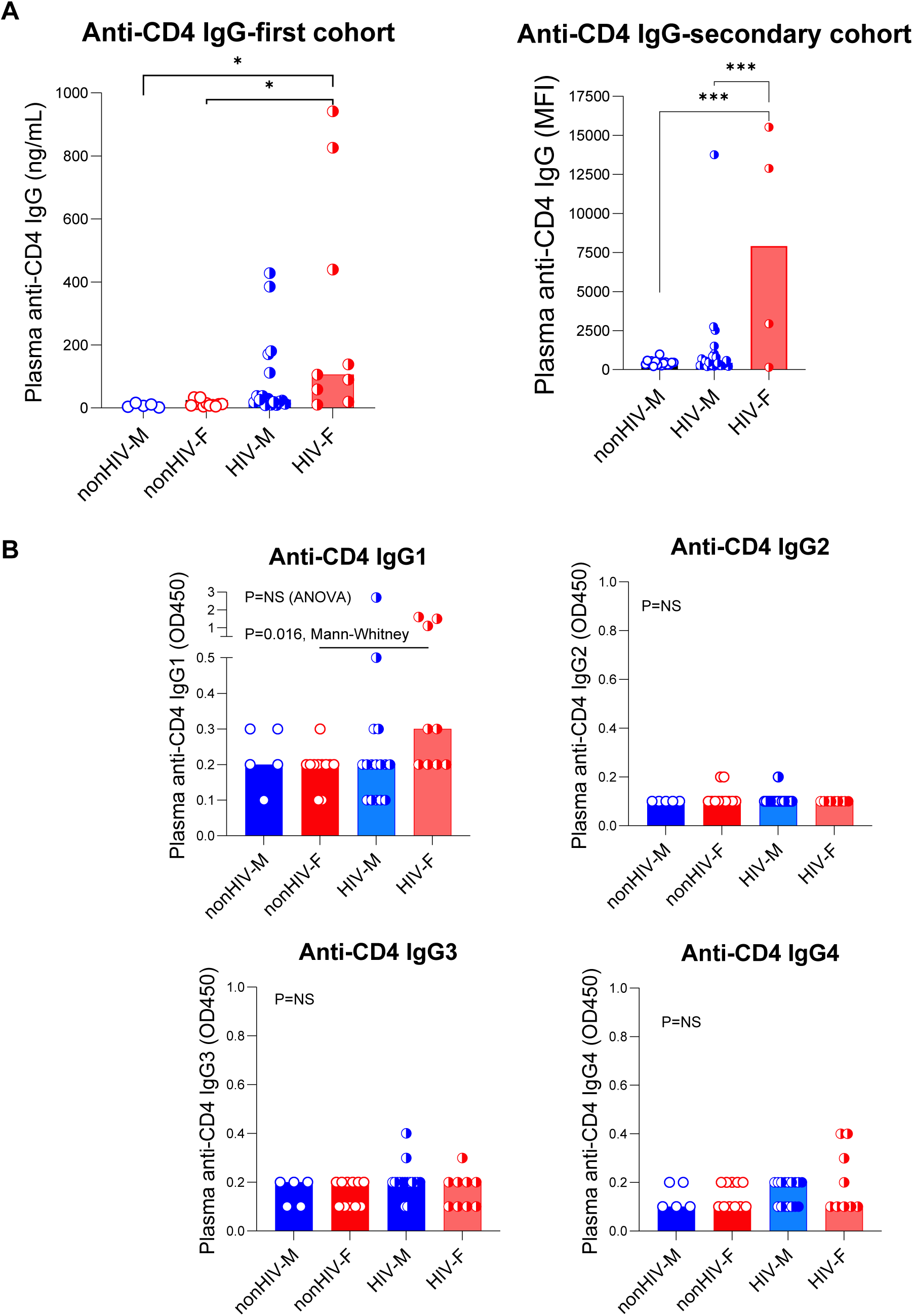
Increased anti-CD4 autoantibody production in PWH on suppressive ART in females but not males. (A) Plasma levels of total anti-CD4 IgG in non-HIV males (non-HIV-M, blue circles), non-HIV females (non-HIV-F, red circles), HIV+ males on suppressive ART (HIV-M, blue squares), and HIV+ females on suppressive ART (HIV-F, red squares). The second cohort (UCSF) had three groups non-HIV males, HIV+ males, and HIV+ females. Data are shown as individual points with median and interquartile range. Statistical significance was determined by one-way ANOVA with post-hoc tests; *p < 0.05. (B) Plasma levels of anti-CD4 IgG subclasses (IgG1, IgG2, IgG3, IgG4) across the same four groups. For IgG1, significance was assessed by one-way ANOVA (P=NS) and Mann-Whitney test between HIV-M and HIV-F (P=0.016). All other subclasses showed no significant differences (P=NS, ANOVA). Data is presented as individual points with median and interquartile range. OD450, optical density at 450 nm.

### Plasma levels of IgGs anti-CD8, anti-ANA, and anti-dsDNA, and sCD4 were similar across the group and lacked correlations with anti-CD4 IgG

To investigate anti-CD4 IgG is unique in HIV, we evaluated other autoantibodies and self-antigen sCD4 for anti-CD4 IgG. Anti-CD8 IgG, anti-dsDNA IgG, ANA IgG, and sCD4 showed no group differences except higher ANA in HIV-M vs. non-HIV-M (*p* = 0.008) and non-HIV-F vs. non-HIV-M (*p* = 0.004, Mann-Whitney U tests; Figure 2A) and higher anti-dsDNA IgG non-HIV-F vs. non-HIV-M (*p* = 0.04, Mann-Whitney U test; Figure 2A). No correlations existed between levels of anti-CD4 IgG and anti-CD8, ANA, anti-dsDNA, or sCD4 (Figure 2B). However, ANA IgG was directly correlated with anti-CD8 IgG in PWH (*r* = 0.41, *p* = 0.04; Figure 2C) and marginally correlated with anti-dsDNA IgG in PWH (*r* =0.34, *p* = 0.08; Figure 2C). These results indicate that anti-CD4 autoimmunity is unique in PWH on ART.

**Figure 2.**
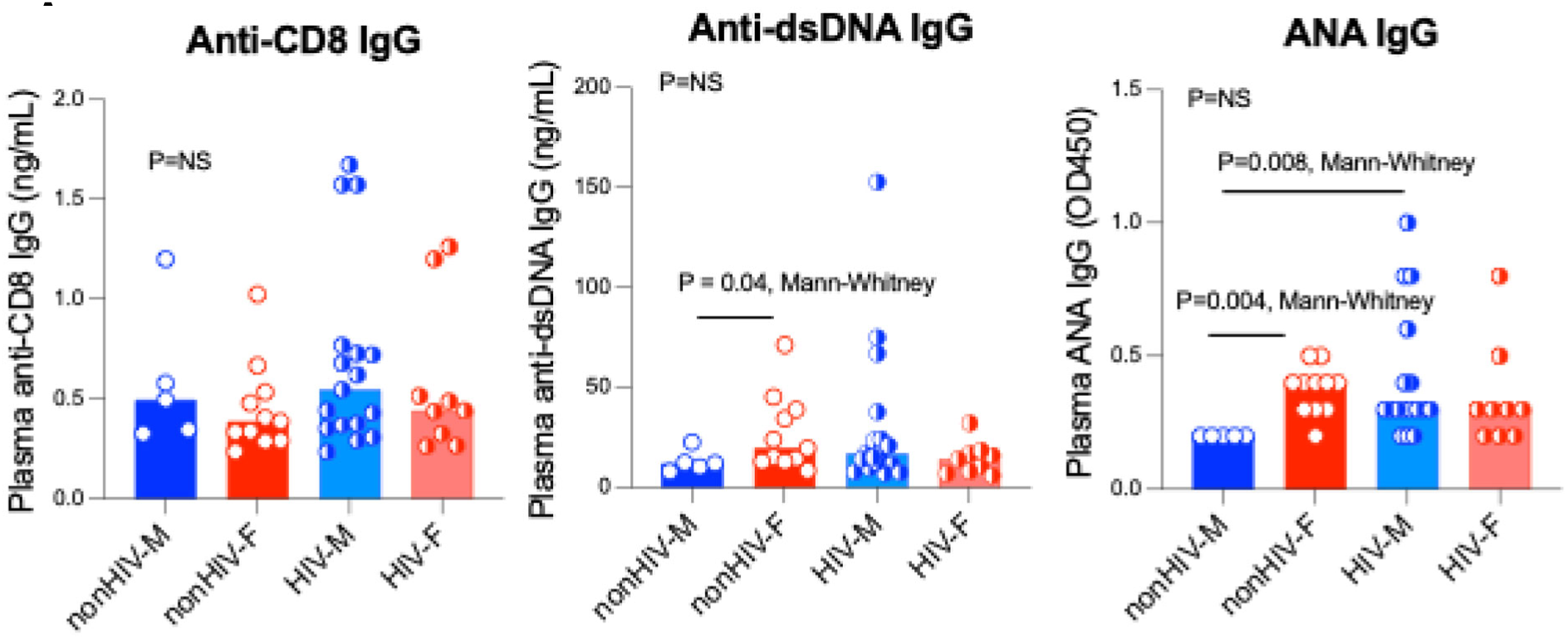
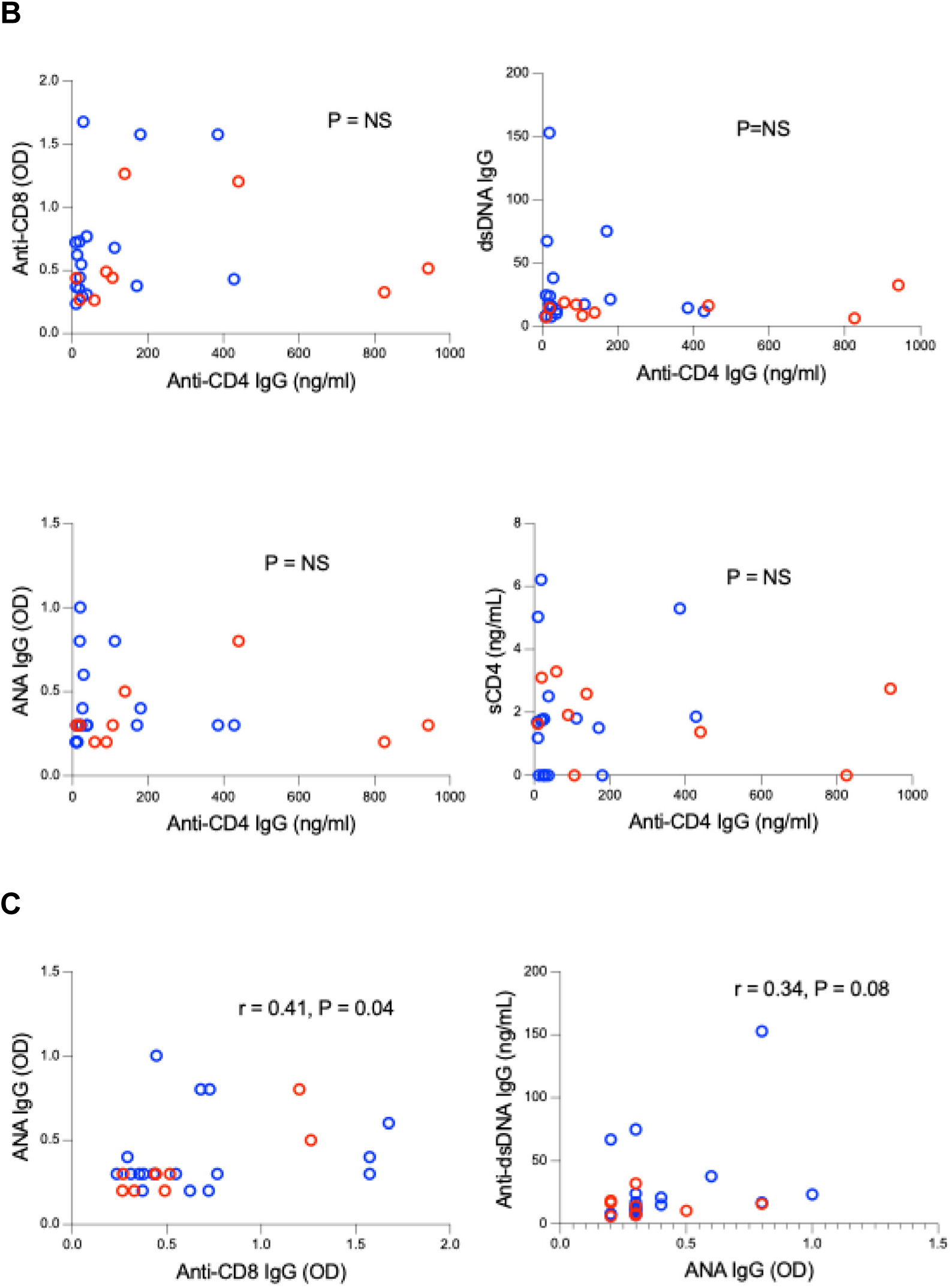
Sex-specific elevation of anti-CD4 IgG in PWH on suppressive ART is not associated with anti-dsDNA IgG, ANA IgG, or soluble CD4. (A) Plasma levels of anti-CD8 IgG (left), anti-dsDNA IgG (middle), and ANA IgG (right) in non-HIV males (non-HIV-M, blue circles), non-HIV females (non-HIV-F, red circles), HIV+ males on suppressive ART (HIV-M, blue squares), and HIV+ females on suppressive ART (HIV-F, red squares). Anti-CD8 IgG showed no significant differences across groups (P=NS, ANOVA). Anti-dsDNA IgG was significantly higher in HIV-F vs. HIV-M (P=0.04, Mann-Whitney); ANA IgG was higher in HIV-F vs. HIV-M (P=0.008, Mann-Whitney) and vs. non-HIV-M (P=0.004, Mann-Whitney). Data are shown as individual points with median and interquartile range. (B) Correlation scatter plots of anti-CD4 IgG (ng/mL) with anti-CD8 IgG (OD), anti-dsDNA IgG (ng/mL), and ANA IgG (OD). No significant correlations were observed (P=NS for all). Correlation of plasma soluble CD4 (sCD4, ng/mL) with anti-CD4 IgG (ng/mL); no significant association (P=NS). (C) Correlation of ANA IgG (OD) with anti-CD4 IgG (ng/mL) in HIV+ individuals (r=0.41, P=0.04, left) and anti-dsDNA IgG (ng/mL) with ANA IgG (OD) (r=0.34, P=0.08, right). Data points represent individual participants; blue = males, red = females. OD450, optical density at 450 nm.

### Inverse correlation between anti-CD4 IgG and blood level of progesterone in HIV+ females

Previous studies have shown that estrogen plays a role in autoantibody production and the pathogenesis of autoimmune disease (15). We next evaluated blood levels of three sex hormones and their correlations with anti-CD4 IgG (Figure 3A-3B). No significant differences in progesterone levels across the four groups (Kruskal–Wallis test, P = NS); no overall significant differences in 17β-Estradiol (Kruskal–Wallis test, P = NS), although HIV+ females showed a trend toward higher levels than non-HIV females. Markedly lower testosterone levels in both female groups compared with both male groups (P < 0.0001, Kruskal–Wallis with Dunn’s multiple comparisons test); no significant HIV-associated differences within the same sex (Figure 3A). Despite the marked female-predominant elevation of anti-CD4 IgG, circulating concentrations of estradiol and testosterone do not correlate with anti-CD4 IgG levels within each sex (Figure 3B). Notably, an inverse relationship was determined between progesterone and anti-CD4 IgG in HIV+ females (Figure 3B). These data confirm the expected physiological sex differences in circulating sex hormone levels, especially testosterone, and demonstrate that long-term virologic suppression on ART does not substantially alter sex hormone profiles in either males or females.

**Figure 3.**
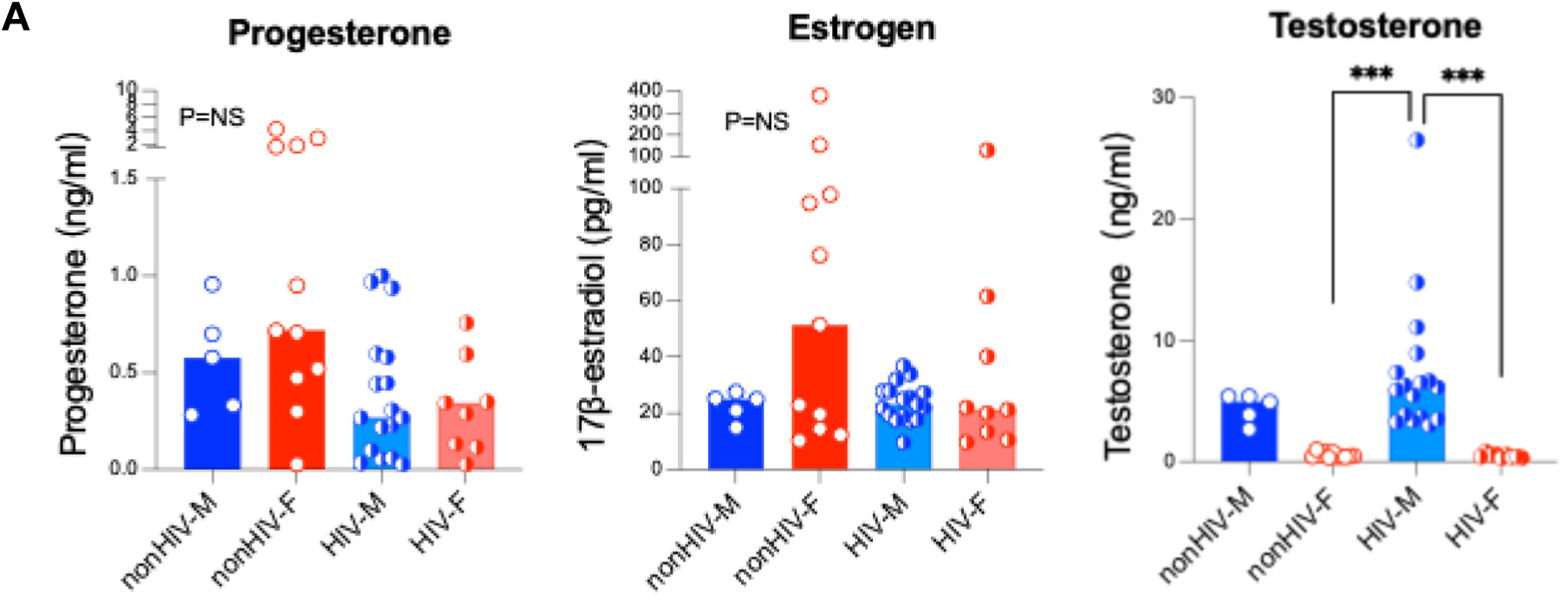
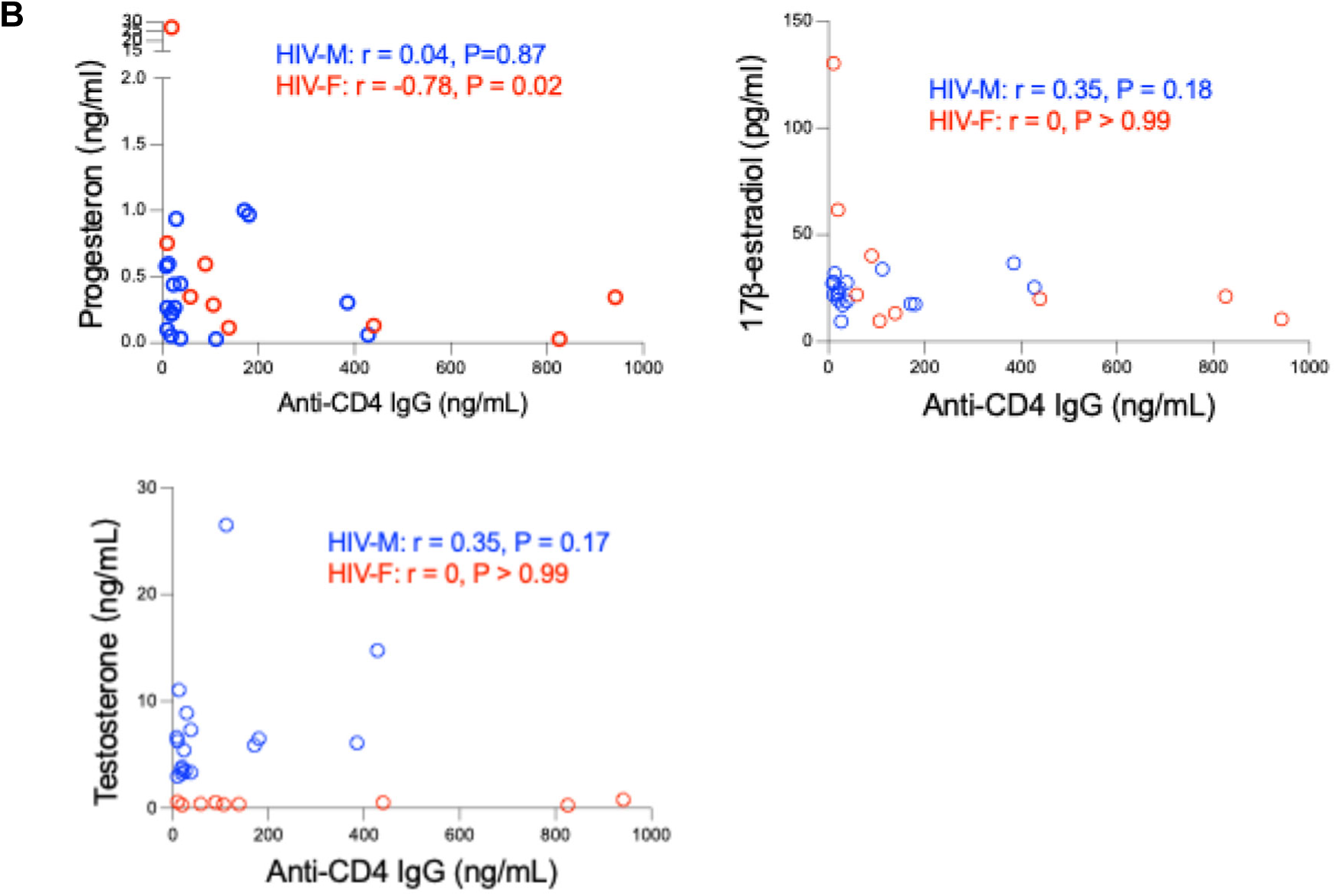
Plasma progesterone levels were inversely correlated with anti-CD4 IgG in HIV+ females. (A) Plasma concentrations of progesterone (left), 17β-estradiol (middle), and testosterone (right) were measured using ELISA in the four study groups. (B) Correlations between anti-CD4 IgG and each sex hormone level. ANOVA and Spearman correlation tests.

### Microbial translocation markers and correlation with anti-CD4 IgG

sCD14 was markedly elevated in both HIV+ males (2.0 ug/mL, 1.5-2.3) and females (2.38 ug/mL, 1.5-2.5) versus non-HIV males (1.45 ug/mL, 1.2-1.6) and non-HIV females (1.43 ug/mL, 1.3-1.5) (\*\**p* < 0.001; Figure 4A). LBP was higher in HIV-F (18.1 μg/mL, 16.6-24.8) versus non-HIV-F (15.3 μg/mL, 13.9-17.2; *p* = 0.016). LPS was also increased in HIV-F (20.5 pg/mL, 16-26.2) versus non-HIV-F (13.1 pg/mL, 10.4-13.9) (*p* < 0.05). All three markers showed female-specific increases in PWH. Among the three markers of microbial translocation, sCD14, LBP, and LPS, only sCD14 correlated positively with anti-CD4 IgG in PWH (*r* = 0.33, *p* = 0.03; Figure 4B), and LBP correlated marginally with anti-CD4 IgG (*r* = 0.35, *p* = 0.08; Figure 4B).

**Figure 4.**
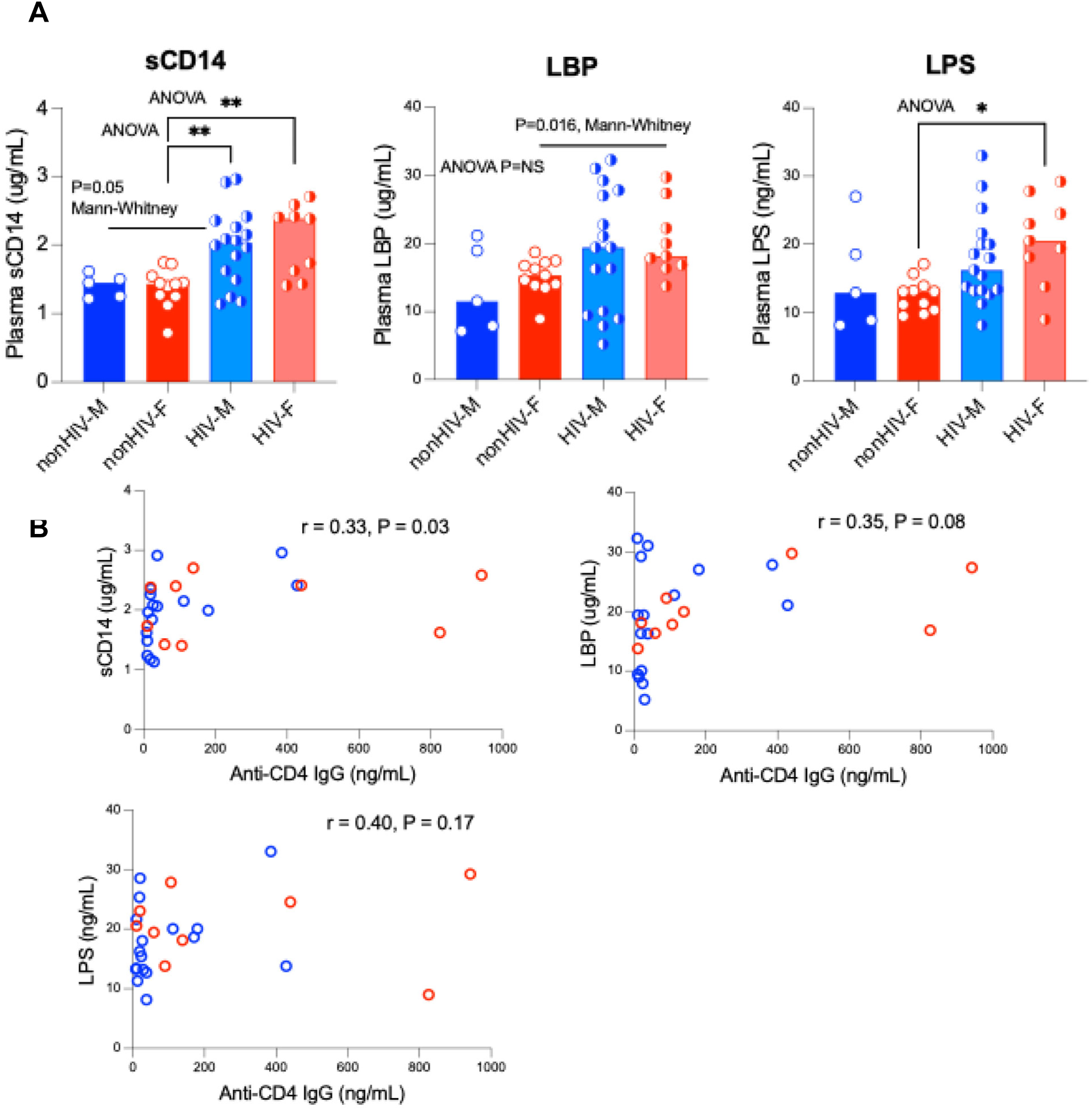
Plasma sCD14 is elevated in people with HIV on suppressive ART irrespective of sex, while LBP and LPS show sex-specific patterns; only sCD14 correlates with anti-CD4 IgG. (A) Plasma levels of soluble CD14 (sCD14, left), lipopolysaccharide-binding protein (LBP, middle), and lipopolysaccharide (LPS, right) in non-HIV males (non-HIV-M, blue circles), non-HIV females (non-HIV-F, red circles), HIV+ males on suppressive ART (HIV-M, blue squares), and HIV+ females on suppressive ART (HIV-F, red squares). sCD14 was significantly higher in both HIV-M and HIV-F vs. non-HIV groups (P<0.005, Mann-Whitney; **P<0.01, ***P<0.001, ANOVA). LBP showed no overall difference by ANOVA (P=NS) but was higher in HIV-F vs. HIV-M (P=0.016, Mann-Whitney). LPS was higher in HIV-F vs. non-HIV-M (P<0.05, ANOVA with post-hoc). Data are individual points with a median. (B) Correlations of anti-CD4 IgG (ng/mL) with sCD14 (r=0.33, P=0.03), LBP (r=0.35, P=0.08), and LPS (r=0.40, P=0.17) in HIV+ individuals. Only sCD14 showed a significant positive correlation. Blue = males, red = females.

### Inverse correlation between anti-CD4 IgG and CD4+ T cell counts

Anti-CD4 IgG correlated negatively with absolute CD4+ T cell count (*r* = −0.59, *p* = 0.002; Figure 5A) but not CD4% or CD4/CD8 ratio (Figure 5B-5C).

**Figure 5.**
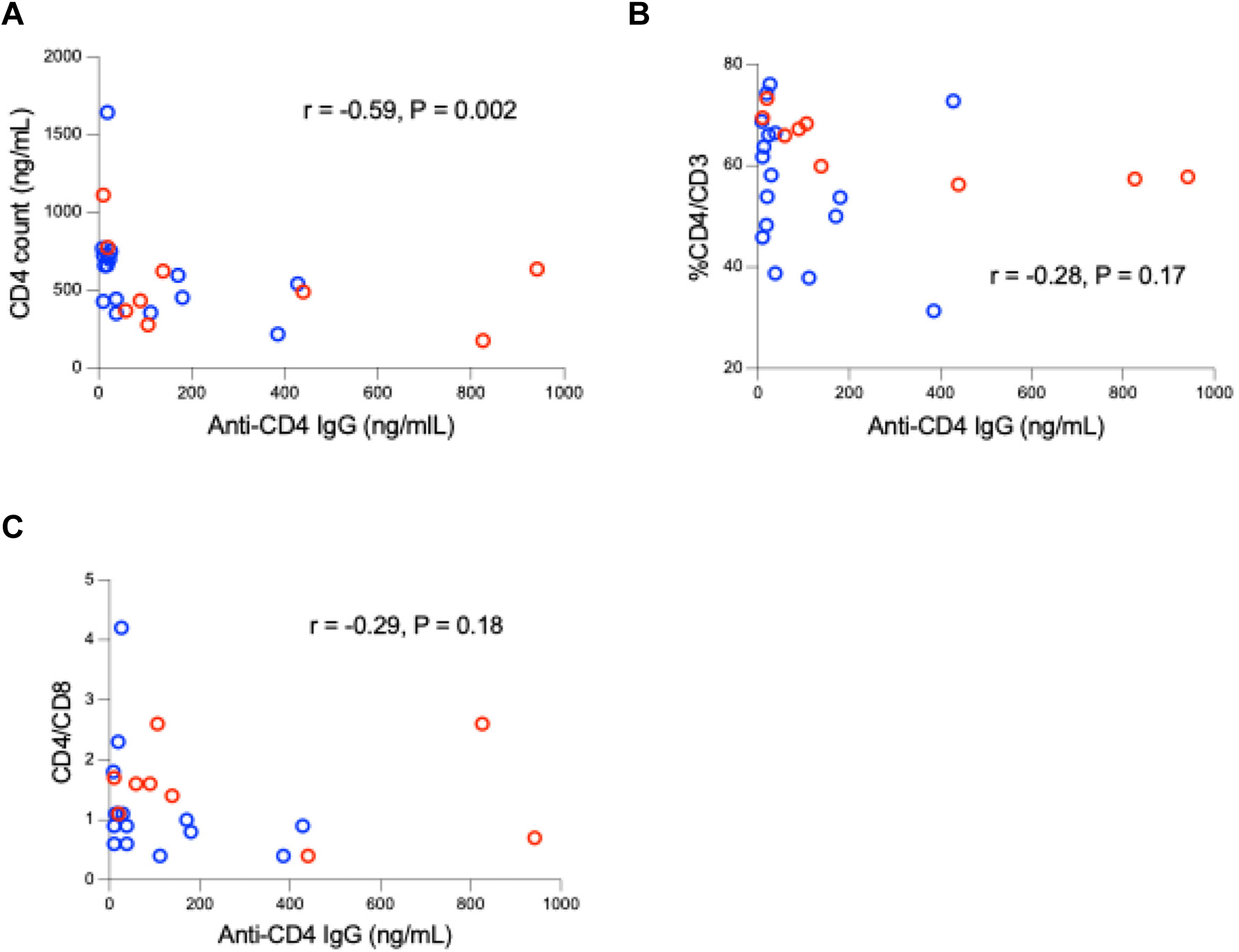
Anti-CD4 IgG levels in people with HIV on suppressive ART correlate with CD4 count but not with %CD4 or CD4/CD8 ratio. Correlation between plasma anti-CD4 IgG (ng/mL) and absolute CD4 count (cells/µL) (A), percentages of CD4 versus CD3 (B), and CD4/CD8 ratio (C) in PWH on suppressive ART. Blue circles = males, red circles = females. Data represents individual participants.

## DISCUSSION

This study establishes anti-CD4 IgG as a previously unrecognized sex-dimorphic autoimmune phenotype in virologically suppressed PWH on ART. The marked female predominance, IgG1 restriction, specificity for CD4 over other self-antigens (versus CD8, ANA, or dsDNA), the absence of IgA and IgM, and the inverse correlation with CD4+ T cell count suggest a targeted, class-switched humoral response that links to treatment outcomes.

In 2017, we first determined that autoimmunity contributes to the outcomes of antiretroviral therapy (success vs. failure) in infectious disease HIV without induction of autoimmune disease (1); later, this concept was well-accepted as shown in studies related to anti-type I IFN autoimmunity in COVID-19 severity from other colleagues and us (2). Several lines of evidence support a causal role for anti-CD4 IgG in blunted immune recovery. First, we and others have previously demonstrated that anti-CD4 antibodies can directly kill CD4+ T cells via ADCC and complement-dependent mechanisms in vitro and ex vivo (1, 16). Second, the strong inverse correlation with absolute CD4 count (r = −0.59) aligns with recent reports linking higher anti-CD4 IgG to immunologic non-response during early ART initiation. Third, the absence of correlation with CD4% or CD4/CD8 ratio suggests the effect is specific to absolute CD4+ T-cell numbers rather than proportional changes driven by CD8 expansion.

HIV+ females experience an elevated risk of premature ovarian reserve depletion and dysregulation of the hypothalamic–pituitary–gonadal (HPG) axis, even when treated with ART (17, 18). The underlying mechanisms remain incompletely understood and are probably multifactorial, involving direct effects of the virus, persistent immune activation and chronic inflammation, lifestyle factors (e.g., smoking), and possibly contributions from

ART itself (19, 20). In the present study, HIV+ females showed a trend toward lower plasma progesterone levels compared with non-HIV females, and progesterone concentrations were inversely correlated with anti-CD4 IgG titers in the HIV+ female group. This relative hypoprogesteronemia aligns with growing evidence of accelerated ovarian aging and HPG-axis dysfunction in HIV+ females (19).

Progesterone exerts potent immunomodulatory effects. It promotes expansion of regulatory T cells, suppresses TLR-induced inflammatory cytokine production by monocytes/macrophages, directly inhibits B-cell activation and class-switch recombination (15), and, as we previously demonstrated, helps maintain gut barrier integrity (21). Consequently, reduced progesterone may lift an important physiological restraint on the chronic, microbial translocation-driven inflammatory milieu that persists in HIV+ females on ART. This loss of restraint could, in turn, promote breakdown of B-cell tolerance and drive the pronounced female-predominant anti-CD4 autoimmune response observed in this cohort. Supporting this concept, progesterone supplementation has already shown clinical benefit in pregnant women with HIV receiving protease inhibitor-based ART (20). These findings warrant further clinical trials to directly evaluate progesterone supplementation or selective progesterone receptor modulators as a novel, sex-specific therapeutic approach to mitigate autoimmune B-cell activation and enhance CD4+ T-cell recovery in women living with HIV.

The female predominance is particularly striking and biologically plausible. Estrogen enhances TLR7/8 signaling in B cells and plasmacytoid dendritic cells, promotes B-cell survival, and lowers the threshold for B-cell activation and class-switch recombination (22). These effects are amplified in the chronic inflammatory milieu of treated HIV infection. The X chromosome encodes multiple immune genes (e.g., TLR7, CD40LG, FOXP3) that escape X-inactivation, providing a potential genetic basis for stronger female humoral responses (23). Microbial products such as LPS and bacterial DNA are potent TLR ligands that can break tolerance via bystander activation or B-cell receptor (BCR) editing; the female-biased elevations in LBP and LPS we observed suggest greater gut barrier dysfunction or heightened innate sensing in women with HIV, which may preferentially drive loss of B-cell tolerance to CD4.

The strong association between anti-CD4 IgG and sCD14 (a cleaved TLR4 co-receptor shed by activated monocytes) mechanistically links this autoantibody response to persistent monocyte activation driven by microbial translocation. Chronic low-level LPS exposure is known to induce polyclonal B-cell activation and autoantibody production in other settings (e.g., systemic lupus erythematosus, primary biliary cholangitis) (24). Our data suggests a similar pathway operates in treated HIV, but with unusual specificity for CD4. This specificity may reflect BCR editing through HIV gp120-bound CD4 (25) or preferential presentation of CD4-derived peptides during chronic immune activation, both of which deserve further investigation.

The paradoxical finding of increased antigen-specific IgG avidity in HIV+ women, despite equivalent or lower total IgG titers in chronic infection (7), further highlights dysregulated but hyper-efficient humoral immunity in females with HIV. Sustained germinal center activity driven by persistent immune activation may favor somatic hypermutation and affinity maturation, resulting in higher-quality antibodies. This “quality over quantity” phenotype has been previously reported for influenza and HIV-specific responses in women (7) and may represent an evolutionary trade-off that confers short-term protection against pathogens at the cost of long-term autoimmunity.

Limitations include the relatively small female sample size (particularly in the second cohort), cross-sectional design, and lack of information on birth control pills and pre- or post-menopausal stages. Longitudinal studies are needed to determine whether anti-CD4 IgG levels precede or follow poor CD4 recovery, and whether they fluctuate with interventions that improve gut integrity. Although anti-CD4 IgGs from some PWH induce CD4+ T-cell death via ADCC, pre-ART levels predict post-ART immune recovery, and B-cell depletion improves CD4+ T-cell reconstitution (8), additional studies, such as comparing anti-CD4 monoclonal antibody function between sexes, cloning CD4-specific B cells, or performing passive-transfer experiments in humanized mice, are needed to confirm pathogenicity.

Clinically, these findings have several implications. First, anti-CD4 IgG may serve as a biomarker to identify PWH at risk for incomplete CD4 recovery, particularly women. Second, therapeutic strategies that reduce microbial translocation (e.g., rifaximin, sevelamer, probiotics, anti-LPS monoclonal antibodies) or directly target pathogenic B-cell responses (e.g., belimumab, daratumumab, BTK inhibitors) could be assessed for their ability to lower anti-CD4 IgG and improve immune reconstitution. Third, progesterone and sex should be considered in the design and interpretation of trials targeting autoimmunity in HIV (i.e., progesterone supplements).

In conclusion, we identify a female-predominant anti-CD4 IgG autoimmune response in treated HIV infection that is linked to low progesterone, microbial translocation, and clinically associated with impaired CD4+ T-cell reconstitution. These findings highlight an under-appreciated intersection of sex, microbial translocation, and autoimmunity in HIV persistence and suggest new therapeutic avenues to improve immune recovery in women.

## AUTHOR CONTRIBUTIONS

D.J., K.S. I.S., A.H., and Z.L., performed experiments. D.J. and Y.Q. wrote the manuscript. Z.W., W. X., D.J., S.S., and W.J. analyzed data. E.C., D. C., L.K., J.E.M., W.S., T.S., S.B., S.F., A.M., and W.J. revised the manuscript.

## ACKOWLEDGEMENTS

This work was supported by grants from the National Institutes of Health R01DA059854 (Jiang), R01DA059538 (Jiang), R01 NS094067 (Price), and the Ralph H. Johnson VA Medical Center Merit Review Award Number I01CX002422 (Jiang) from the United States (U.S.) Department of Veterans Affairs Office of Research and Development (CSR&D) Service, as well as the Swedish government and the county councils (ALF agreement ALFGBG-965885, Gisslen), the Swedish Research Council (#2021-06545, Gisslen), and by SciLifeLab from the Knut and Alice Wallenberg Foundation (2020.0182 & 2020.0241, Gisslen).

## DATA AVAILABILITY STATEMENT

Data are available on request.

---

**Supplemental Figure 1.**
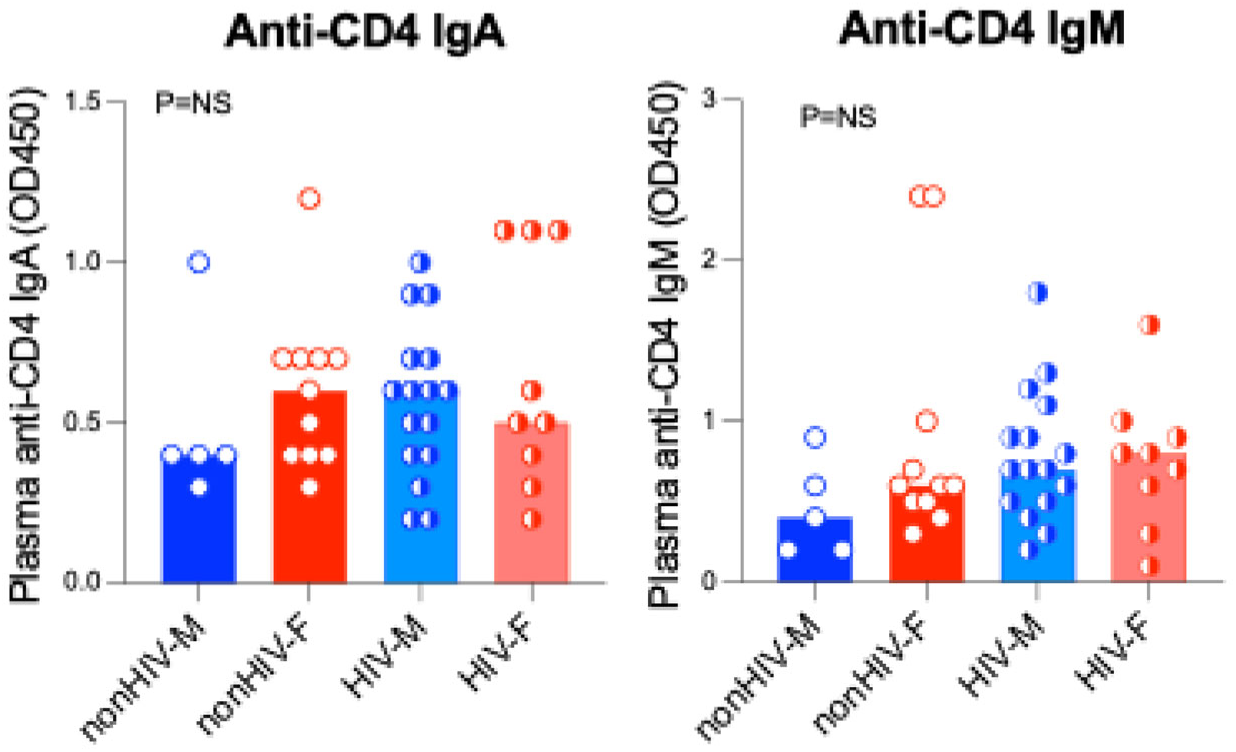
No detectable anti-CD4 IgA or IgM in plasma from PWH on suppressive ART or uninfected controls. (A) Plasma anti-CD4 IgA levels in non-HIV males (non-HIV-M, blue circles), non-HIV females (non-HIV-F, red circles), HIV+ males on suppressive ART (HIV-M, blue squares), and HIV+ females on suppressive ART (HIV-F, red squares). No significant differences across groups (P=NS, ANOVA). (B) Plasma anti-CD4 IgM levels across the same four groups. No significant differences were observed (P=NS, ANOVA). Data are shown as individual points with median and interquartile range. OD450, optical density at 450 nm.

## Reference

1. Luo Z, Li Z, Martin L, Wan Z, Meissner EG, Espinosa E, Wu H, Yu X, Fu P, Julia Westerink MA, Kilby JM, Wu J, Huang L, Heath SL, Li Z, Jiang W. 2017. Pathological Role of Anti-CD4 Antibodies in HIV-Infected Immunologic Nonresponders Receiving Virus-Suppressive Antiretroviral Therapy. J Infect Dis 216:82–91.

2. Bastard P, Rosen LB, Zhang Q, Michailidis E, Hoffmann HH, Zhang Y, Dorgham K, Philippot Q, Rosain J, Beziat V, Manry J, Shaw E, Haljasmagi L, Peterson P, Lorenzo L, Bizien L, Trouillet-Assant S, Dobbs K, de Jesus AA, Belot A, Kallaste A, Catherinot E, Tandjaoui-Lambiotte Y, Le Pen J, Kerner G, Bigio B, Seeleuthner Y, Yang R, Bolze A, Spaan AN, Delmonte OM, Abers MS, Aiuti A, Casari G, Lampasona V, Piemonti L, Ciceri F, Bilguvar K, Lifton RP, Vasse M, Smadja DM, Migaud M, Hadjadj J, Terrier B, Duffy D, Quintana-Murci L, van de Beek D, Roussel L, Vinh DC, Tangye SG, et al. 2020. Autoantibodies against type I IFNs in patients with life-threatening COVID-19. Science 370.

3. Gazzola L, Tincati C, Bellistri GM, Monforte A, Marchetti G. 2009. The absence of CD4+ T cell count recovery despite receipt of virologically suppressive highly active antiretroviral therapy: clinical risk, immunological gaps, and therapeutic options. Clin Infect Dis 48:328–37.

4. Lisco A, Wong CS, Lage SL, Levy I, Brophy J, Lennox J, Manion M, Anderson MV, Mejia Y, Grivas C, Mystakelis H, Burbelo PD, Perez-Diez A, Rupert A, Martens CA, Anzick SL, Morse C, Chan S, Deleage C, Sereti I. 2019. Identification of rare HIV-1-infected patients with extreme CD4+ T cell decline despite ART-mediated viral suppression. JCI Insight 4.

5. Bowler SA, Premeaux TA, Ratzan L, Friday C, Gianella S, Landay AL, Ndhlovu LC, NWCS ACTG. 2025. Plasma anti-CD4 IgG levels are associated with poor immune recovery in people with HIV initiating antiretroviral therapy. AIDS 39:208–210.

6. Migliore L, Nicoli V, Stoccoro A. 2021. Gender Specific Differences in Disease Susceptibility: The Role of Epigenetics. Biomedicines 9.

7. Luo Z, Ogunrinde E, Li M, Zhang L, Martin L, Zhou Z, Hu Z, Zhang T, Li Z, Zhang J, Su B, Zhang T, Wu H, Ma L, Liao G, Eckard AR, Westerink MAJ, Heath SL, Jiang W. 2019. Increased influenza-specific antibody avidity in HIV-infected women compared with HIV-infected men on antiretroviral therapy. AIDS 33:33–44.

8. Epling BP, Lisco A, Manion M, Laidlaw E, Galindo F, Anderson M, Roby G, Sheikh V, Migueles SA, Poole A, Perez-Diez A, Liu X, Rao VK, Burbelo PD, Sereti I. 2024. Impact of Anti-CD4 Autoantibodies on Immune Reconstitution in People With Advanced Human Immunodeficiency Virus. Clin Infect Dis doi:10.1093/cid/ciae562.

9. Moran JA, Turner SR, Marsden MD. 2022. Contribution of Sex Differences to HIV Immunology, Pathogenesis, and Cure Approaches. Front Immunol 13:905773.

10. Botey-Bataller J, van Unen N, Blaauw M, Vos W, van Eekeren L, Vadaq N, Matzaraki V, Verbon A, Groenendijk AL, Dos Santos JC, Cleophas MCP, Stalenhoef JE, Berrevoets MAH, Jiang X, Gupta MK, Nguyen N, Xu CJ, Joosten LAB, Netea MG, van der Ven A, Li Y. 2025. Genetic and molecular landscape of comorbidities in people living with HIV. Nat Med 31:3350–3359.

11. Stehle JR, Jr., Leng X, Kitzman DW, Nicklas BJ, Kritchevsky SB, High KP. 2012. Lipopolysaccharide-binding protein, a surrogate marker of microbial translocation, is associated with physical function in healthy older adults. J Gerontol A Biol Sci Med Sci 67:1212–8.

12. Brenchley JM, Price DA, Schacker TW, Asher TE, Silvestri G, Rao S, Kazzaz Z, Bornstein E, Lambotte O, Altmann D, Blazar BR, Rodriguez B, Teixeira-Johnson L, Landay A, Martin JN, Hecht FM, Picker LJ, Lederman MM, Deeks SG, Douek DC. 2006. Microbial translocation is a cause of systemic immune activation in chronic HIV infection. Nat Med 12:1365–71.

13. Cheng D, Luo Z, Fu X, Stephenson S, Di Germanio C, Norris PJ, Fuchs D, Ndhlovu LC, Li QZ, Zetterberg H, Gisslen M, Price RW, Peng S, Jiang W. 2022. Elevated Cerebrospinal Fluid Anti-CD4 Autoantibody Levels in HIV Associate with Neuroinflammation. Microbiol Spectr doi:10.1128/spectrum.01975-21: e0197521.

14. Luo Z, Li M, Wu Y, Meng Z, Martin L, Zhang L, Ogunrinde E, Zhou Z, Qin S, Wan Z, Westerink MAJ, Warth S, Liu H, Jin P, Stroncek D, Li QZ, Wang E, Wu X, Heath SL, Li Z, Alekseyenko AV, Jiang W. 2019. Systemic translocation of Staphylococcus drives autoantibody production in HIV disease. Microbiome 7:25.

15. Moulton VR. 2018. Sex Hormones in Acquired Immunity and Autoimmune Disease. Front Immunol 9:2279.

16. Luo Z, Jiang W. 2024. A protocol for anti-CD4 IgG antibody purification using plasma samples from people with HIV and antibody-mediated cytotoxicity. MethodsX 12:102698.

17. Guzha BT, Mateveke B, Mubata H, Chapupu T, Dondo V, Chirehwa M, Tshikosi R, Chipato T, Chirenje ZM. 2025. Assessment of the impact of HIV infection on the hypothalamic-pituitary-ovarian axis and pubertal development among adolescent girls at a tertiary centre in Zimbabwe: a cross-sectional study. BMC Endocr Disord 25:16.

18. Yalamanchi S, Dobs A, Greenblatt RM. 2014. Gonadal function and reproductive health in women with human immunodeficiency virus infection. Endocrinol Metab Clin North Am 43:731–41.

19. Unachukwu CN, Uchenna DI, Young EE. 2009. Endocrine and metabolic disorders associated with human immune deficiency virus infection. West Afr J Med 28:3–9.

20. Siou K, Walmsley SL, Murphy KE, Raboud J, Loutfy M, Yudin MH, Silverman M, Ladhani NN, Serghides L. 2016. Progesterone supplementation for HIV-positive pregnant women on protease inhibitor-based antiretroviral regimens (the ProSPAR study): a study protocol for a pilot randomized controlled trial. Pilot Feasibility Stud 2:49.

21. Zhou Z, Bian C, Luo Z, Guille C, Ogunrinde E, Wu J, Zhao M, Fitting S, Kamen DL, Oates JC, Gilkeson G, Jiang W. 2019. Progesterone decreases gut permeability through upregulating occludin expression in primary human gut tissues and Caco-2 cells. Sci Rep 9:8367.

22. Young NA, Wu LC, Burd CJ, Friedman AK, Kaffenberger BH, Rajaram MV, Schlesinger LS, James H, Shupnik MA, Jarjour WN. 2014. Estrogen modulation of endosome-associated toll-like receptor 8: an IFNalpha-independent mechanism of sex-bias in systemic lupus erythematosus. Clin Immunol 151:66–77.

23. Youness A, Miquel CH, Guery JC. 2021. Escape from X Chromosome Inactivation and the Female Predominance in Autoimmune Diseases. Int J Mol Sci 22.

24. Hang L, Slack JH, Amundson C, Izui S, Theofilopoulos AN, Dixon FJ. 1983. Induction of murine autoimmune disease by chronic polyclonal B cell activation. J Exp Med 157:874–83.

25. Yoon V, Fridkis-Hareli M, Munisamy S, Lee J, Anastasiades D, Stevceva L. 2010. The GP120 molecule of HIV-1 and its interaction with T cells. Curr Med Chem 17:741–9.

